# Cortex-Wide Cellular Imaging in Freely Locomoting Mice Using Cortex Camera Array Microscope (CortexCAM)

**DOI:** 10.64898/2026.02.11.705445

**Authors:** Arun Cherkkil, Zoey Viavattine, Vamsy Kota, Kapil Saxena, Daniel Surinach, Ibrahim Oladepo, Jill Juneau, Jia Hu, Skylar M.L. Fausner, Eunsong Ko, Shubham Panchal, Sanjana Srivatsa, Malachi R. Lehman, Roarke Horstmeyer, Suhasa B Kodandaramaiah

## Abstract

Understanding single-cell neuronal activity across multiple brain regions in the context of ethologically relevant behaviors is a major goal in systems neuroscience. We have engineered the Cortex Camera Array Microscope (CortexCAM), integrating four miniaturized fluorescence imaging microscopes to simultaneously capture cellular activity from contiguous fields of view spanning over 48 mm^2^ of the dorsal cortex. The CortexCAM is capable of imaging > 9000 individual neurons across much of the primary and secondary motor, somatosensory, visual, retrosplenial, and association cortices across both hemispheres of the dorsal cortex. The compact nature of the CortexCAM allows integration into a passive mechanical gantry system to form the mobile CortexCAM. The mobile CortexCAM allows volitional control of the animal’s translational motion (x, y) and rotational motion (yaw) in physical behavior arenas. We then use the mobile CortexCAM to perform cortex-wide cellular resolution imaging in freely locomoting mice performing alternating choice tasks, as well as during social interactions. Thus the CortexCAM allows studying cortex-wide cellular dynamics in behaviors that cannot be achieved in headfixed settings.

## INTRODUCTION

In recent years, considerable efforts have been made to understand how widespread regions of the cortex coordinate their neural activity to mediate complex behaviors. These studies have revealed key insights into how attention, task engagement, motor activity and arousal all recruit cortical circuits globally and how activity across the cortex is coordinated to mediate these behaviors. These studies have been facilitated primarily by two recent advances. First, the ability to optically access large regions of the cortex using curved^1–6^ or flat cranial windows^7^, and second, the advent of imaging systems capable of imaging large field of views (FOVs) at high resolution^8–12^.

While these studies have revealed key insights into how disparate regions of the cortex are engaged during behavior, these benchtop devices are typically many orders of magnitude larger in size than the mice they are imaging, requiring head-fixation. Headfixed imaging devices are limited in their ability to study complex behaviors requiring self-motion, such as spatial navigation, and sensory motor integration, such as eye-head coupling. Such experiments do not recapitulate the wide repertoire of behaviors seen in freely behaving mice. While several attempts have been made to develop wide-field-of-view miniaturized microscopes that can be head-borne by freely behaving animals^13–19^, miniaturization reduces the overall device performance. Currently, no miniaturized device has the ability to image the entire cortex at the signal-to-noise ratio and spatial temporal resolution comparable to a benchtop imaging system.

Here, we present an ultra-compact imaging system (Cortex Camera Array Microscope or ‘CortexCAM’) capable of imaging the entire dorsal cortex of mice (>48mm^2^ of imaging area) at cellular resolution. The CortexCAM utilizes an optical design framework that integrates four miniaturized fluorescence microscopes to simultaneously capture cellular calcium activity from overlapping fields of views of the dorsal cortex in mice. This approach gives an ability to image a 48mm^2^ wide area of the mouse cortex at single-cell resolution. Furthermore, we engineered a mobile version of the CortexCAM (mobile CortexCAM) that can image widespread neuronal cortical populations in freely locomoting mice exploring physical behavioral arenas. We show that the CortexCAM can simultaneously image over 9000 neurons in freely locomoting mice. We demonstrate the utility of the CortexCAM by surveying whole cortex cellular dynamics in mice performing a naturalistic alternating choice task.

## RESULTS

### CortexCAM hardware

The CortexCAM, fuses four miniaturized fluorescence imaging microscopes into one integrated imaging system (**Fig. 1 a,b**). Each microscope is reminiscent of designs employed by miniaturized microscopes^20,18,21,16^, with two key differences. First each imaging microscope incorporates a collimated light source (**Fig. 1c**) that delivers a much higher light output (90 ± 5 mW/mm^2^) than typical light emitting diodes (LEDs) integrated into a miniaturized microscope. Second, a molded aspheric lens (**Fig. 1c**) is used instead of gradient refractive index (GRIN lens)^21^ or compound lenslet assemblies^20^ that allows wide-FOV imaging while also providing a long working distance (15.2 mm). The microlens allows near 1:1 imaging of the imaging field and maps 6.50 mm x 5.86 mm area of the imaging plane to 1140 by 900 pixels in the CMOS sensor. Each pixel has an area of 2.9 µm^2^. This compact design allows four imaging microscopes to be fused together into a single housing that is shown in **Fig. 1c**. Each microscope has an independent focus mechanism that allows the CMOS sensor position to be adjusted to bring the imaging plane into focus. The cameras are positioned to image planes with slight angular offsets (29 degrees between two posterior microscopes, 30 degrees between two anterior microscopes, and 15 degrees between each pair of anterior-posterior microscopes). A corresponding convex polyhedral double glass coverslip implant (**Fig. 1b**, also see Methods) creates four imaging planes normal to the optical imaging axes of each of the four microscopes of the CortexCAM^22^.

**Figure 1.**
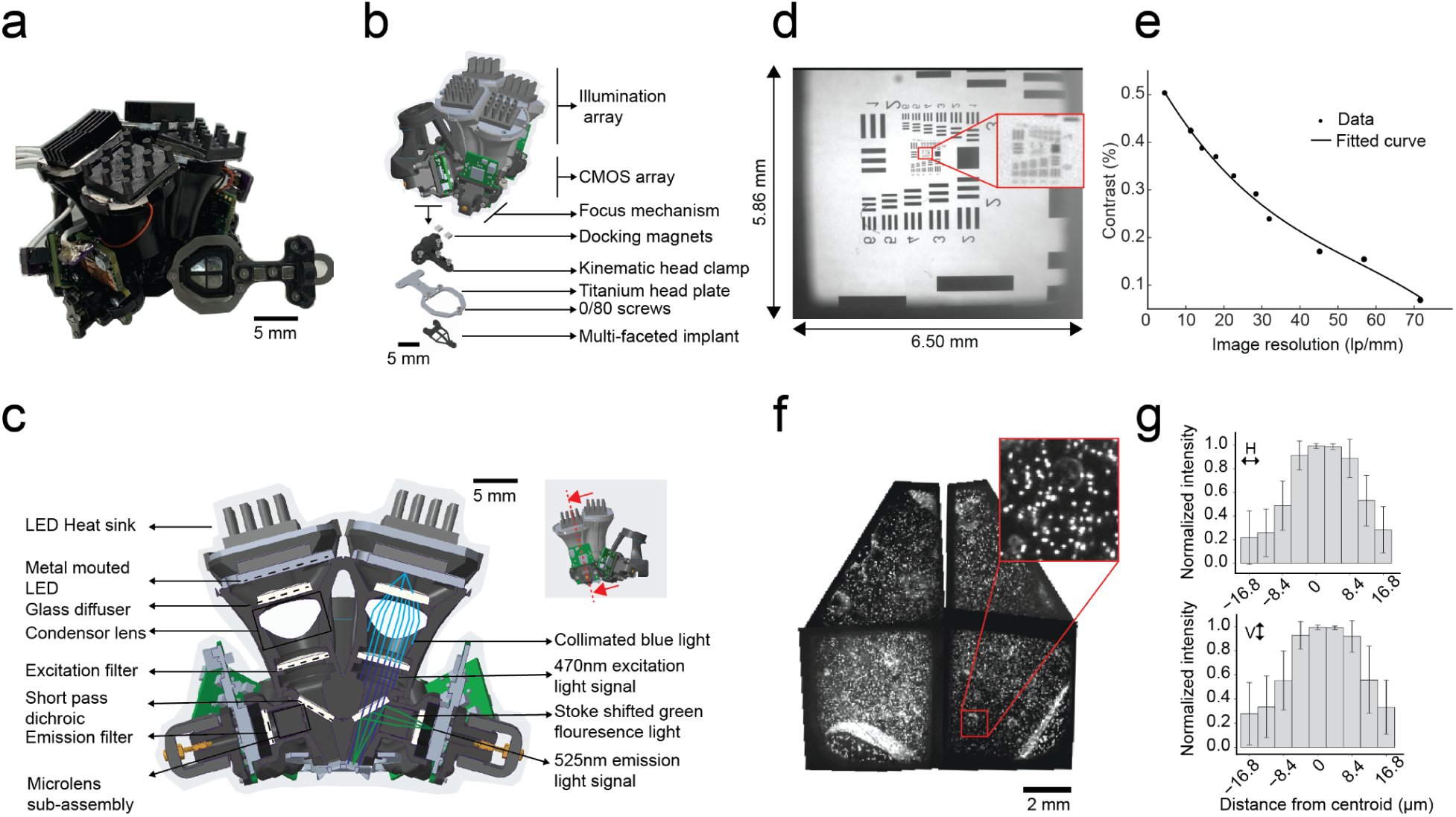
The CortexCAM: A compact microscope for cortex-wide cellular imaging. a) Photograph of the CortexCAM and the corresponding polyhedral coverglass implant b) Isometric CAD view of the CortexCAM with arrows demarcating the excitation, emission, focusing sub-assemblies. Bottom: polyhedral coverglass implant sub assembly. c) Cross-sectional view of the CortexCAM: Right half cross-section shows the ray-traces for excitation and emission light paths. The left half cross-section shows the individual opto-electronic components. d) United States Airforce (USAF) test target imaged using a single imaging module of the CortexCAM. Inset expands and shows group 6 and 7. e) Modulation Transfer Function (MTF) curve approximated from experimental values obtained using micro-objective lenses used in each CortexCAM imaging module. f) 10 µm fluorescence microbeads in a phantom implant imaged using the CortexCAM. Inset shows a small region of microbeads imaged from one imaging field of view (FOV). g) Average light intensity variation across individual microbeads indicated in the inset (f).

The parallelized microscope architecture results in an array of four microscopes, each independently imaging contiguous FOVs at cellular resolution, that cumulatively give a coverage of >48mm^2^ area of the dorsal cortex. The overall footprint of the CortexCAM is approximately 55 x 52 x 45 mm^3^, and results in a device weight of 49.1 g including the tethers. The two posterior microscopes image through cranial windows, each covering an area of 4.10 x 4.10 mm^2^, encompassing higher order visual cortex, somatosensory cortex and retrosplenical cortex brain regions in both hemispheres. Correspondingly, each of the anterior microscopes images 8.405 mm^2^ area comprising motor and prefrontal cortices (**Fig. 2a**) Cumulatively, >48 mm^2^ of the dorsal cortex can be imaged. Each microscope within the CortexCAM has a working distance of 15.2 mm.

**Figure 2.**
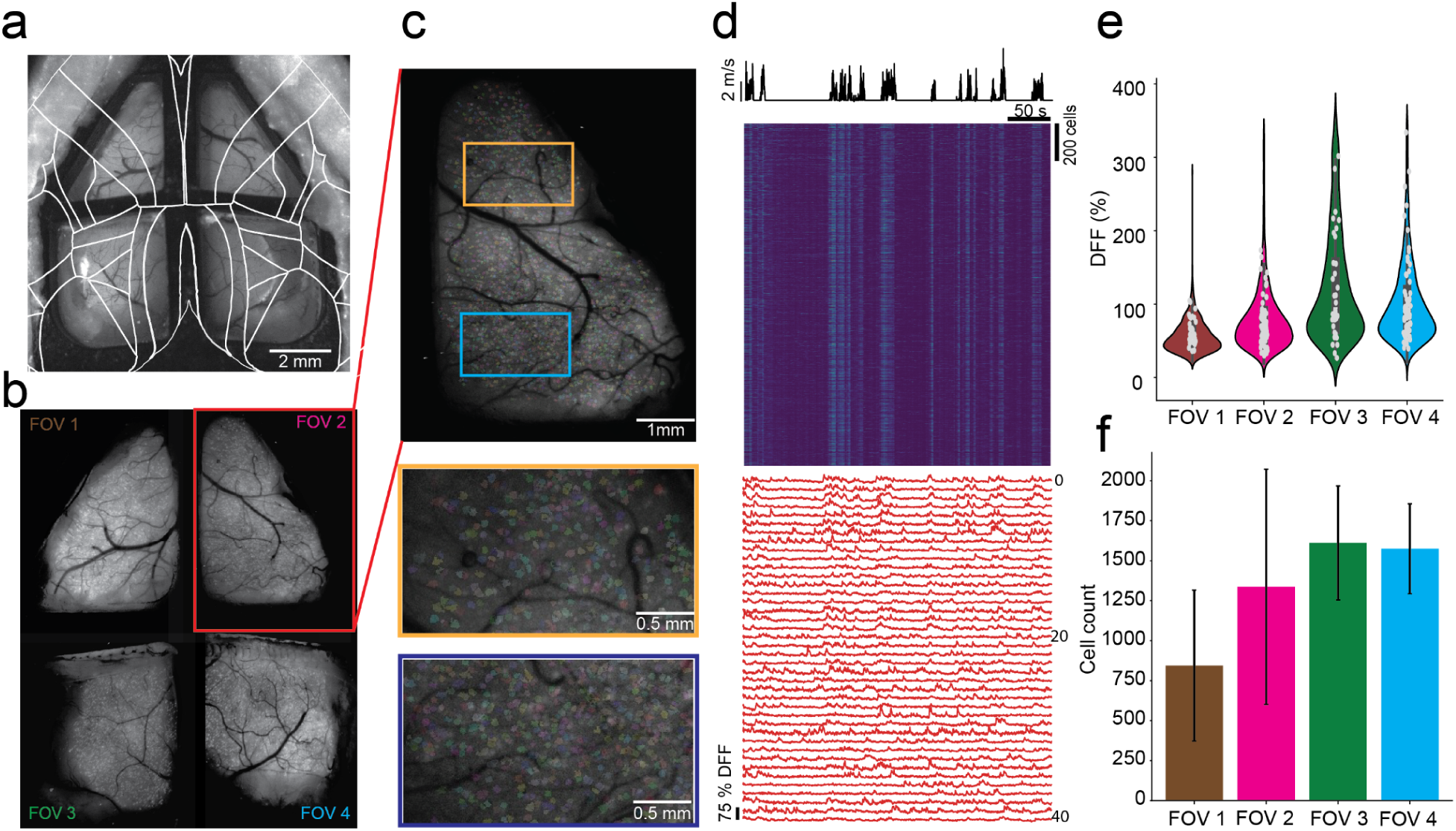
Cortex-wide cellular resolution imaging in head-fixed mouse. a) Widefield image of a Cux2-CRE-ERT2 x Ai162^3^ mouse implanted with the convex polyhedral double glass coverslip implant. White lines indicate an overlaid brain atlas^59^. b) A composite max projection image of 4 simultaneously imaged FOVs using the CortexCAM. c) Expanded view of FOV 3 showing footprints of the extracted cell regions (top) and two further expansions on the two rectangular insets marked as orange and blue in the first expanded view (bottom). d) *Top:* Pseudo color heatmaps of normalized DFF traces for all neurons imaged in FOV 3 (top) along with velocity of the locomotion on the treadmill. *Bottom:* representative Ca2+ DFF times series of 20 randomly selected neurons. e) Distribution of peak DFF (<95^th^ percentile) of all neurons imaged in each FOV (n = 3 mice). f) Number of neurons recorded in each FOV (n = 3 mice).

A third distinction is a kinemantic clamping mechanism (**Fig. 1b**) built into the bottom surface of the CortexCAM. The clamping mechanism allows mice, implanted with a convex polyhedral double glass coverslip cranial window, to be docked directly to the compact microscope unlike headfixation to a post over a treadmill.

### CortexCAM Optical Characterization

Each excitation and emission light paths within the CortexCAM is independent, and is designed to be identical to the other paths with minimal variations in imaging resolution and field of view. Each microscope employs a plastic molded aspheric pinhole lens assembly that is sourced from a commercially available portable microscope with a measured numerical aperture of 0.053 NA and an effective focal length of 3.75 mm (See **Methods**).

The aspheric microlens has a fixed focal length and could provide varying magnifications by changing the relative distance of the specimen and the CMOS sensor from the lens. The microlens comes in a molded plastic housing that has a diameter of 5.1 mm and length of 6 mm. It also incorporates a pinhole opening on the specimen facing side that facilitates rejection of out-of-focus light from the imaging light path.

To optimize for the maximum field of view without compromising resolution, we performed focus tests using USAF resolution targets at different working distances and back focal lengths for the lens and sensor combination (**Supplementary Info. 1a**). In order to image from the largest optical window in the implant (two posterior windows) we needed to achieve a field of view of 4.1 x 4.1 mm with minimal distortion. The plastic lenses typically have best resolution at the center of FOV, with spherical aberrations introduced in the periphery^13^. In the final configuration, we were able to achieve an imaging FOV of 74.21 mm^2^ at full pitch resolution of 13.9 μm at a working distance of 15.1 mm (**Fig. 1d)**. This allowed us to reduce the effective occupancy of each imaging cranial window to <45% across the FOV, thereby minimizing peripheral aberrations.

To further quantify the imaging quality of the system, we measured the contrast (%) across different spatial frequencies. We performed a quadratic fit over all experimental data points and observed a contrast level > 20% at a spatial frequency of 71.8 lp/mm, which corresponds to our application specific critical resolution of 13.9 μm. With a sparse expression of GCamp6s in layers 2/3 of the transgenic mice, coupled with image processing algorithms for signal extraction, we were able to discriminate activity of single neurons at this resolution^12,13^.

Next we imaged 10μm fluorescent microbeads to check for aberration in the corner regions of the brain windows (**Fig. 1f)**. We measured the average light intensity variation across a corner sub-region of a brain window and reported the Full Width Half Maximum (FWHM) across both horizontal and vertical direction to be 16.8μm. This shows minimal aberration in the corner regions. The excitation light source for the CortexCAM is a custom light torch sub-assembly that uniformly illuminates (**Supplementary Info. 1b**) each brain window with clear blue excitation light with a sharp cut-off at 491.2 nm and a light power delivering capacity of 95mW/mm^2^ at the object side aperture of the scope (**Supplementary Info. 1c**).

We performed *in vitro* testing of a fully assembled single microscope module by imaging a 400μm coronal brain slice from a mouse expressing GCamp6s^3,23^ sparsely in layers 2/3 excitatory neurons using CortexCAM (**Supplementary Info. 1d**) and verified that individual neuronal regions sparsely expressing GCamp6 could be distinguished.

### Cortex-wide headfixed imaging using the CortexCAM

We next evaluated the performance of the CortexCAM in headfixed imaging experiments. The convex polyhedral double glass coverslip implants were implanted on Cux2/Cre-ERT2 mice crossed with Ai162(TIT2L-GC6s-ICL-tTA2) mice^3^. Each illumination module provided 19 ± 2 mW/mm^2^ of 470nm filtered blue light illumination to the brain. Mice were docked to the bottom mounting feature of the CortexCAM while running on a custom-built treadmill. We were able to clearly identify thousands of neurons in each FOV (**Fig. 2b,c, Supplementary Video 1, 2**). **Fig. 2d** shows the calcium activities of all cells recorded from a single mouse over a 5 minute trial while ambulatory. The running speed was captured using an encoder attached to the wheel. We saw clear increases in Ca^2+^ activity throughout the dorsal cortex corresponding to locomotion. On average, we found that neurons had peak DFF value of 50 ± 15.70 % (6152 neurons) and this was consistent across all four FOVs that we imaged. We were able to record from 844 ± 471 and 1337 ± 735 neurons (n = 3) in the anterior brain windows covering prefrontal, somatosensory, and sections of the motor-cortical regions bilaterally. Additionally, in the posterior fields of view covering the motor cortex, high order visual and the retrosplenial cortical regions bilaterally, we were able to identify 1998 ± 412 and 1574 ± 281 neurons (n=3). The slight differences in the number of neurons recorded in the anterior and posterior regions of view can be attributed to the overall differences in imaging areas. The anterior windows typically cover an imaging area of 16 mm², whereas the posterior windows cover an area of 32 mm². Further, there can be variations in the overall expression of GCaMP in cells in these regions^3,13,24^.

### Mobile CortexCAM for imaging in minimally restrained mice locomoting physical behavioral arenas

Previously, several efforts have focused on developing highly miniaturized imaging devices that can image the brain in freely behaving animals^21,25–28^. These miniaturized devices allow recording of neural activity in a wide repertoire of behaviors assays. Typically, miniaturized imaging systems are designed to weigh less than 4 g, so as to allow them to be head-borne by animals freely exploring behavioral arenas^29^. While the CortexCAM is considerably more compact than most typical bench-top microscopes having comparable imaging performance, the weight of the CortexCAM does not allow for head-borne deployment. However, mice, which have a higher strength/weight ratio than rats, have the ability to exert forces sufficient to lift, displace, or move objects with masses greater than their own^30^. Here we realized a novel mechanical gantry platform to suspend the CortexCAM, allowing it to translate and rotate freely in a horizontal plane, which can then be maneuvered volitionally by mice to explore physical spaces (**Fig. 3a**). This mobile version of the CortexCAM can be incorporated over behavioral arenas as large as 80 cm x 50 cm. The CortexCAM is essentially suspended overhead the behavior arena using linear rails that can translate in both X and Y, as well as a rotational bearing that allows the CortexCAM to rotate about a yaw axis.

**Figure 3.**
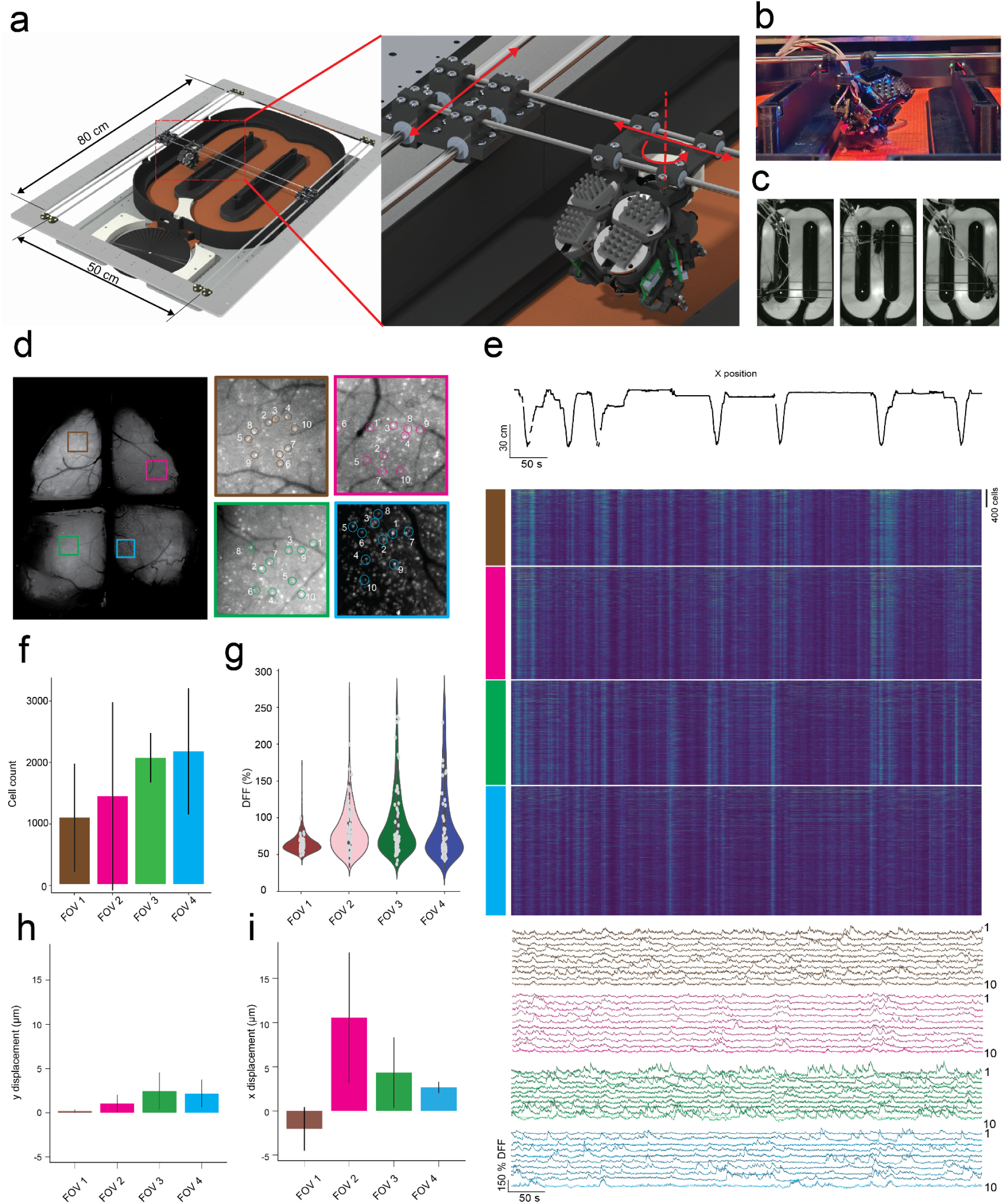
Mobile CortexCAM for cortex-wide cellular imaging in freely locomoting mice. a) CAD schematic of the mobile CortexCAM integrated over a figure 8 maze behavior arena. The expanded view shows a zoomed in image of the CortexCAM with the arrows representing the x, y and yaw degrees of freedom. b) Photograph of a mouse navigating the 8-maze behavior arena while docked to the mobile CortexCAM. c) Series of images captured during free exploration of the behavioral arena. d) A composite max projection image of 4 simultaneously imaged FOVs by the mobile CortexCAM from a mouse freely exploring the 8 maze arena (left). Expanded views of small sub-FOVs as marked in the composite image. White circles indicate 10 randomly selected neurons whose Ca^2+^ DFF time cells are highlighted in (e). e) *Top:* The x-displacement of the mouse in the figure 8 maze area during a single 5 minutes trial. *Bottom:* Psuedo-color heat maps of the deconvolved Ca^2+^ DFF time series of all neurons imaged in the 4 FOVs. *Bottom:* The Ca^2+^ DFF traces for all 40 cells highlighted in (d). f) Number of neurons recorded in each FOV (n = 3 mice). g) Distribution of peak DFF (<95^th^ percentile) of all neurons imaged in each FOV (n = 3 mice). h) Rigid motion displacement in the y direction for each of the FOVs (n =3 mice). i) Rigid motion displacement in the x direction for each of the FOVs (n = 3 mice).

Mice readily acclimatized to being docked to the mobile CortexCAM and maneuvered the imaging device around the entire behavioral arena in 6-7 training sessions with each session lasting more than 10 minutes. (**Fig. 3b, c, Supplementary Video 3**). In a social interaction arena where a stranger mouse was introduced in a confined cage, a mouse tethered to the mobile CortexCAM (subject mouse) readily interacted socially with the stranger mouse (**Supplementary Video 4**). The mice were constrained in their ability to rotate their heads about the pitch and roll axes. First, we evaluated the effect of docking mice to the mechanical gantry platform on the locomotion of the mice. We evaluated both the distance moved and the speed of the mice exploring figure 8 maze arena when docked to the gantry without a scope (effective weight of 7 g), an intermediate (weight of 28 g), and a weight of 52 g (equivalent to the fully assembled CortexCAM), and compared that to locomotion observed when mice were docked to including all the wiring tethers that may cause additional torsion. Despite the varying loads and torsions, mice moved similar distances during a 10 minute trial regardless of the load they were maneuvering (41.89 ± 4.02 m at 7 g load, 47.01 ± 5.43 m at 28 g load, 49.52 ± 3.95 m at 52 g load and 51.30 ± 5.76 m with CortexCAM, n = 3 mice, **Supplementary Info. 3a, b**) There were no statistically significant differences in the median speed during the trials (6.790 ± 0.693 cm/s at 7 g load, 8.12 ± 0.90 cm/s at 28 g load, 8.26 ± 0.64 cm/s at 52 g load and 9.44 ± 0.83 cm/s with CortexCAM, n = 3 mice, **Supplementary Info. 3c, d**). The same mice while not tethered to the gantry platform moved a total distance of 67.24 ± 6.87 m (n = 3), not significantly higher than the tethered mice (p > 0.08, Friedman test). The median speed of locomotion was also comparable to the speeds obtained while animals were docked to the CortexCAM (9.93 ± 1.10 cm/s when untethered and freely behaving, 9.44 ± 0.83 cm/s when tethered to mobile CortexCAM, p > 0.1, Friedman test). Thus, the mobile CortexCAM, in principle, allows high-resolution, wide-field, cortex-wide imaging at a performance level similar to bench-top microscopes, but in minimally restrained animals that are behaving naturalistically in a 2D behavioral arena.

### Whole cortex cellular imaging in freely locomoting mice

We performed cellular resolution imaging across the entire dorsal cortex using the mobile CortexCAM while animals were exploring a figure 8 maze arena (**Fig. 3d-i**). **Fig. 3d** shows the maximum intensity projection from a single imaging trial where a total of 9491 cells were recorded across the four imaging fields of view. We were able to record neural activities as mice made multiple traverses across the entire figure 8 maze arena. During movement, DFF traces showed a distinct increase in activity at the onset of movement. This is similar to several other works that have shown that movement onset results in broad activation of the cortex^31–34^. On average, we were able to record from 4017 ± 1093 cells in both the posterior fields of view, encompassing most of the retrospinal, posterior somatosensory, and the visual cortices across both hemispheres (n = 3 mice, **Fig. 3f**). In the two anterior windows encompassing the primary and secondary motor cortices, and parts of the somatosensory and prefrontal cortices, we were able to record 2473 ± 1719 cells (n = 3 mice). Average peak DFF registered during the trial (**Fig. 3d)** was 57 ± 15.70 % across all the cells recorded (**Fig. 3g**). We next evaluated the stability of imaging as mice traversed the maze by quantifying the degree to which each imaged field of view needed to be corrected for motion artifacts using a rigid body motion correction algorithm^35,36^. On average, in both X and Y directions, rigid body motion correction was limited to less than 10 μm (n=3) in each field of view. There were variations across different fields of view possibly owing to minor optical misalignments or mechanical misalignments of individual cameras. On average, across all fields of view, rigid body motion correction was limited to 1.46 ± 0.91 μm in the X direction and 3.88 ± 4.50 μm in the Y direction (**Fig. 3h, i**). Thus, the mobile CortexCAM can perform stable, high SNR Ca^2+^ imaging 1000s of individual cells across the whole cortex in minimally restrained mice engaging in complex behaviors.

### Cortex-wide encoding of heading direction in an alternating choice task

We imaged the dorsal cortex when mice were trained to perform an alternating choice task^37,38^ while tethered to the mobile CortexCAM. Mice navigated the figure 8 maze arena, traversing through the central decision arm, taking alternating left and right turns to receive a reward in the peripheral arms. Mice performed an average of 32 ± 7.21 trials (**Supplementary Info. 3f)** in each 10-minute experimental session (**Supplementary Info. 3f** and **g**). Although the same mice completed more trials when untethered and freely behaving, the difference was not statistically significant (47 ± 7.55 untethered and freely behaving, 32 ± 7.21 tethered to mobile CortexCAM, n = 3, p>0.07 Friedman test). Freely behaving mice had significantly lower successful choices (46.99 ± 11.72 % freely behaving, 91.59 ± 11.14 % tethered to mobile CortexCAM, n = 3, p<0.04, Friedman test; post-hoc Tukey test: p<0.003).

In contrast to head-fixed imaging systems, mice on the mobile CortexCAM have volitional control over their heading direction and movement in 2D spatial arenas. Mice readily showed a proclivity to explore areas where they received rewards (**Supplementary Info. 3h**), while also demonstrating possible vicarious trial-and-error-type behaviors at the decision-making zone^39^(**Supplementary Info. 3e, Supplementary Video 3**). Mice had the ability to make 180 degree turns and backtrack along the path. Marker-less tracking of mouse behavior during the figure 8 maze task showed that the heading direction of the mice were distributed evenly with slightly higher proclivity for heading direction in either the 0° or the 180° angles owing to the asymmetry in the behavioral arena (**Supplementary Info. 3h)**.

We used markerless tracking^40^ to estimate both the position and the heading direction of the mice as they performed the alternating choice task (**Fig. 4a**). Owing to the asymmetric nature of the figure 8 maze, much of the angles were confined to the azimuth angle of 0° while mice were traversing the two outer arms towards the reward, or 180° when the mice were traversing the center decision making arm(**Fig. 4b**). Overall, there were few instances of mice being oriented in other directions, and these were primarily while the animals were making the decision to turn to either of the outer arms or as they made their way into the central arm once they received the reward. Within the context of the mice performing this alternating choice task, we looked at whether neurons in the cortex were tuned to firing in a particular heading direction. Using statistical analysis (see **Methods**), we found that a fraction of the recorded neurons were significantly more likely to fire in a particular heading direction, and their activities could be sorted as a function of the heading direction of the mice as they performed the alternating choice task (**Fig. 4c and d**).

**Figure 4.**
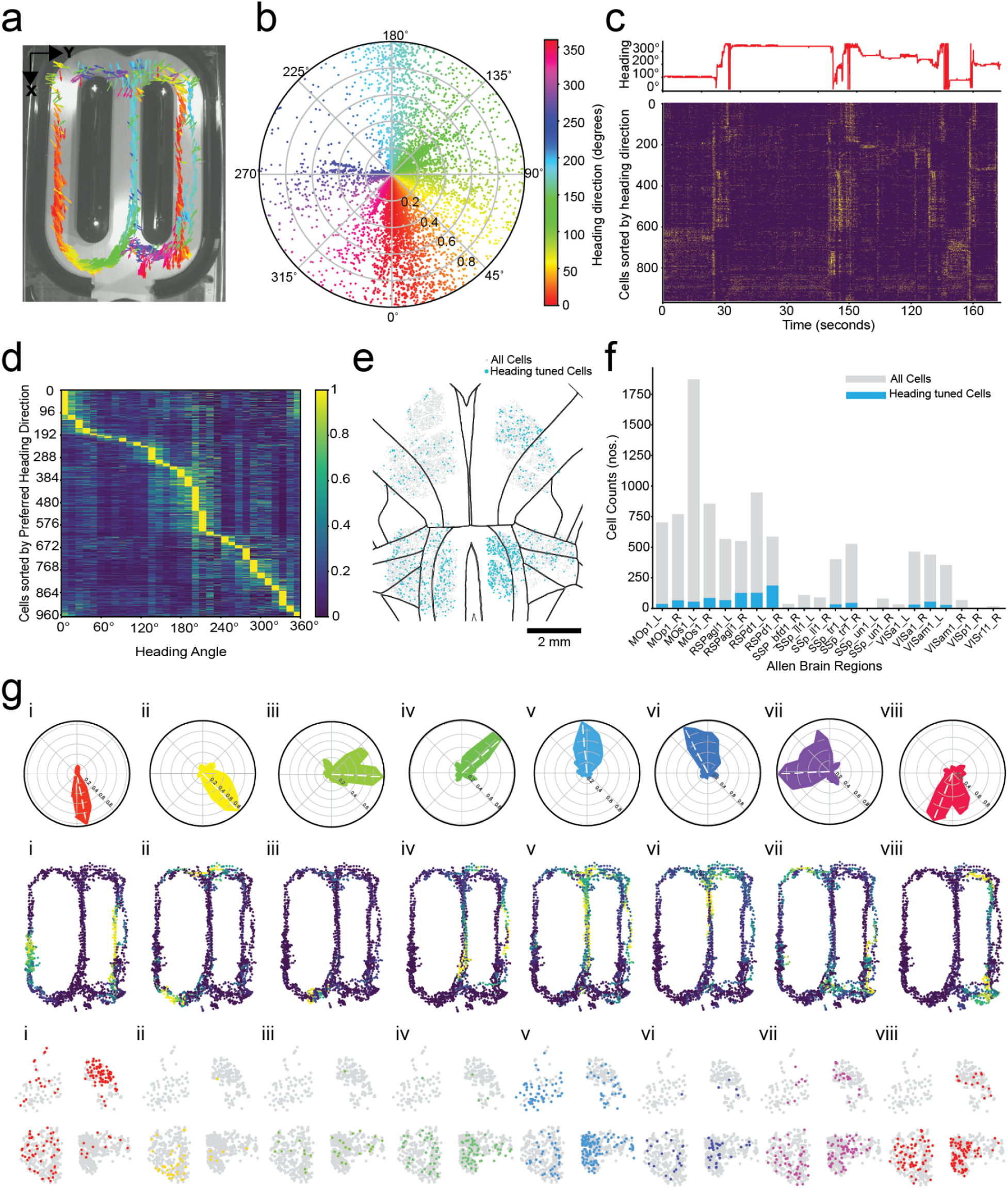
Distributed encoding of heading direction in the dorsal cortex. a) markerless tracking of heading direction and position of a mouse performing alternating choice tasks in the figure 8 maze. b) Polar scatter plot of instantaneous heading direction of the mouse with distance from center along the radial axis indicating the instantaneous normalized speed. c) *Top*: The heading direction of the mouse during a 180 s time interval selected from the full trial shown in **a** and **b**. *Bottom*: Pseudo color heatmap of the deconvolved spike trains for a subset of cells tuned to the heading direction of the mouse, sorted by their preferred heading direction. d) Pseudo-color heatmap indicating the trial averaged normalized tuning strength of the filtered subset of cells tuned to the heading direction of the mouse. Cells are sorted by their preferred heading direction. e) Location of all cells (gray circles) recorded during the session overlaid with cells tuned to the heading direction of the mouse (blue circles). f) The breakdown of number of cells imaged and those encoding heading direction by cortical regions. g) *Top:* The trial averaged normalized tuning curves for 8 representative cells each occupying a different heading direction bin. *Middle:* The deconvolved Ca^2+^ trace for the cells during the whole 10 minute trial. *Bottom:* A scatter plot showing all cell regions (gray circles) overlaid with the cells with preferred heading direction falling within a 45° angle bin centered about the peak activation angle (colored circles) for each cell shown above.

**Fig. 4e** shows the distribution of 9491 cells recorded in a single trial, indicated using gray circles, and blue circles indicating the locations of 968 (10.20%) cells that were found to be tuned to the heading direction of the mouse during the alternate choice task. We found that cells encoding heading directions were distributed throughout the cortex in all recorded regions. However, there were differences in the proportion of cells that were found to have heading direction properties. For instance, we found a higher fraction of cells (19.12%) that were encoding heading direction in the retrosplenial cortex, whereas in the bilateral motor cortices, the fraction was much lower (5.7%) (**Fig. 4f**). We also examined whether there were differences in the distribution of cells encoding a particular direction. **Fig. 4g** shows representative examples of cells that fire in eight different directions (**Fig. 4g.i** - **viii**). Consistent with previous studies, motion towards a reward in a goal-directed fashion resulted in a higher activation of cells in the anterior regions of the cortex^41–43^ (**Fig. 4g.i**). In contrast, motion in the central arm (**Fig. 4g.v**) and decision arms (**Fig. 4g.iii - iv** and **vi-vii**) indicated a higher fraction of cells in the posterior, particularly retrosplenial regions, having a higher density of heading direction cells. Thus, the mobile CortexCAM uniquely enables survey of encoding properties of single cells distributed over very large areas of the cortex otherwise not possible with existing head-fixed imaging systems.

## DISCUSSION

Compared to large benchtop microscopes^12,23,44–48^, the CortexCAM has a highly compact form factor. This compact form factor allows integration into the low-friction sliding support system to realize the mobile CortexCAM that can be volitionally manipulated by mice exploring and locomoting in physical arenas. Moreover, the overall imaging FOV of the CortexCAM is comparable and in some cases supersedes the FOV of these benchtop systems, while also allowing cortex-wide cellular resolution Ca^2+^ imaging during naturalistic behaviors in mice.

Traditionally, large scale neural imaging has been achieved using either large bench-top microscopes where animals are limited to headfixed behaviors, or have been performed using small FOVs using miniaturized microscopes. While head mounted scopes allow complex behaviors to be studied, they also result in increased behavioral variability and add to experimental confounds. In contrast, headfixed experiments in mice allow precise constraints on behaviors, but require significant training and motivation. The mobile CortexCAM draws from the key advantages of both head-fixed and freely behaving imaging systems. Specifically, the docking style and negligible contamination of neural data from motion and edge distortion artifacts makes it easy to adopt this platform for any head-fixed behaviors. The naturalistic behaviors of the mice allow training of behaviors similar to freely behaving assays, with the added constraint of limiting motor behavior to X, Y translation and yaw rotation. Such behaviors can be accomplished while a supraminiaturized (∼50 g) imaging device is integrated. This simple mechanical system could serve as a blueprint for developing new technologies where miniaturization to under 2 g devices^17,20,49^ is not possible or difficult to achieve. This could open up a new design space for compact neural imaging and perturbation devices.

In contrast with headfixed mice behaviors, where self motion is not present, volitional control of heading direction and sensory motor integration allows novel ways for animals to interact with their physical environment. Head fixation results in disruption in eye-head movement coupling. This is essential for mice for shifting gaze during visually guided behaviors^50^. Using the mobile CortexCAM platform, we have overcome this limitation by allowing the mice tethered to the platform to have unrestrained movement in the X,Y and yaw directions while simultaneously recording cortex-wide neural activity.

Finally, the CortexCAM was realized by adapting hardware assembly methods that have been optimized by miniaturized microscopic imaging systems^25,28^. As shown in supplementary video 5, the assembly is relatively straightforward. This significantly reduced the overall expense and dependency on otherwise restrictive specialized benchtop microscopes to achieve comparable imaging results. The data acquisition is compatible with the open-source miniscope ecosystems and can thus be easily incorporated into existing experimental pipelines. In conclusion, we anticipate both the miniaturized realization of the CortexCAM as well as the mechanical gantry system for supporting such a supraminiaturized system to open up a novel design space where new kinds of recording and stimulation devices can be designed for use by systems neuroscientists.

## METHODS

### Convex polyhedral double glass coverslip implant

The structural element of the custom convex polyhedral double glass coverslip implant was 3D printed in black resin using a stereolithography 3D printer (Form2, Formlabs Inc.). The 3D printed bezel partitions the implant into four cavities. Each cavity was inlaid with two layers of custom cut glass coverslips of 170 µm thickness (# 89428-986, VWR Cover glasses No.1). Each cavity consisted of grooves to position the custom glass coverslip pieces. First, a thin layer of optical adhesive (NOA61, Norland Optical Adhesives Inc.) was applied along the grooves. The first glass coverslip piece was carefully placed in the cavity to allow the adhesive to spread uniformly between the two surfaces. A UV light (315-400 nm) torch was used for 60 seconds to cure the adhesive and firmly hold the glass piece in position. A second glass coverslip layer was then placed on top of the first layer. The optical adhesive was gently applied along the edges of the glass coverslip using an insulin syringe needle tip (0.3mL Insulin Syringes, SOL-M™) to allow the adhesive to form a uniform layer between the two glass coverslips through capillary action. The UV torch was then used to cure the adhesive and form a double layered optically transparent cranial window. This process is repeated for the remaining 3 cavities to form the convex polyhedral double glass coverslip implant. A titanium headpost is attached to the cranial window implant using three screws.

The headpost incorporated a CNC machined Delrin clamp at the posterior end. The Delrin clamp included two square shaped neodymium magnets (B221, K & J magnets) of size 1/8 x 1/8 x 1/16 inches to allow for quick headclamping, as done previously^24^.

### The CortexCAM opto-mechanical construction

The CortexCAM device housing is divided into three key sub-assemblies. The top housing combines four independent excitation light paths that allow us to provide uniform illumination across all four fields of view. The middle housing is designed to house four independent emission light-paths, that are deflected by 90 degrees from their respective excitation light paths to simultaneously capture neural activity from all four implant window partitions. The middle housing is further modified to be integrated with the gantry platform using three M2 brass screws (96741A017, McMaster-Carr). Each of the four arms of the middle housing incorporates manually actuated focusing mechanisms that axially translate the CMOS sensor by 0.25 mm per revolution, to make fine focus adjustments.

#### Top excitation light housing module assembly

The top housing is split along the centerline of each of the four light modules to form a single merged middle female half housing and four mating male half housings to allow for precise placement of the optical elements within each of the four excitation light paths. The light module parts were 3D printed in black ABS material(Accura 7820, Protolabs) using a high resolution stereolithography printer (Form3, Formlabs Inc.). Each illumination module consists of a metal mounted blue LED (M470D4, Thorlabs). The LEDs are custom shaped by removing material from the edges of the metal mount using the Electrical Discharge Machining (EDM) process. A mini-finned composite heatsink (04021 heatsink, LEDsupply) is attached to the bare side of the LED metal mount using thermally conductive tape (TCDT1, Thorlabs). Each of the light module additionally comprises an excitation filter (10mm diameter, 1.1mm thickness, ET470/40x, Chroma Technologies) and an aspheric condenser lens (12.7 mm diameter, 0.78 NA, 7.5mm central thickness, ACL12708U, Thorlabs). The excitation filter and the condenser lens were first placed within their corresponding grooves in the middle housing and fixed in place by dabbing small amounts of optical adhesive (NOA61, Norland Optical Adhesive) along the edges of the housing that is in contact with the optical elements, followed by UV light exposure for curing. This process was then repeated for the remaining three illumination sub-modules. Once the middle housing is complete with four pairs of excitation filter and condenser lens, the mating male parts are press-fitted onto the middle section using snaps. A small piece of copper tape (Kraftex copper tape, 1/4 inch, Amazon) is placed along the mating surfaces to prevent light leakage. The LEDs are then positioned in the groves provided along the top surface of the housing and sealed using an optical adhesive (NOA74, Norland Optical Adhesive) (**Supplementary Video 5**).

#### Middle emission light module assembly

The experimenter first placed the scope body, pitched upwards, on a polystyrene petri-dish that is filled with a reusable adhesive putty material. Once the scope is anchored in the putty, the experimenter places a 7 x 5 x 1 mm custom cut short pass dichroic in the groove provision made on the top surface of one of the front-facing arms using rubber tipped tweezers. Prior to placing the dichroic in the housing, the surface is cleared of any dust particles by spraying the surface with compressed air followed by dipping the dichroic in 70 % ethanol solution and finally drying of the coated surfaces using a narrow tip cotton applicator with the contacting tip rolled up with a small cut piece of lens tissue paper (MC-5, Thorlabs). It is important to have the reflective coated surface of the dichroic to be cleared of any debris to avoid scattering of fluorescence signals that are deflected by the dichroic mirror surface to form the final image by passing through the emission light path. The short pass dichroic is oriented at 45 degrees with the incoming light to maximize the filter efficiency. UV-curing optical adhesives (NOA 61, Norland) is used to affix the dichroic by dabbing a small quantity of the adhesive along the corners of the mating surface of the dichroic housing using an insulin syringe tip. This process is repeated for the remaining three arms of the scope body, with the two rear facing arms having groove provisions made for holding a 7 x 7 x 1 mm custom cut dichroic (**Supplementary Video 5**).

In order to design a simple optical framework, we utilized a molded aspheric lens extracted from a portable microscope (MS100, Teslong). The lens is then press-fitted onto a lens holder sub-assembly that is machined out of high performance plastic (Delrin, Protolabs). The lens holder also has a provision on image facing side of the lens to insert a 6*6*1mm custom cut emission filter (ET 535/30m, Chroma Technology Corp). The filter is fixed in place by dabbing small quantities of optical adhesive along the corners of the filter enclosure and UV curing the glue. This subassembly is then inserted to the lens enclosure in the middle housing module. The experimenter further applies small quantities of the optical adhesive along the 4 corners of the mating surfaces to restrict any misalignment due to the clearance fit between the lens holder and the enclosure. A custom laser cut piece of compressible polyurethane foam (93275K113, McMaster-Carr) is attached to the bottom face of the enclosure for the CMOS sensor, with a central circular cutout to allow light to pass through the lens and form an image on the sensor. The CMOS sensor sub-assembly is then carefully placed within the cavity provided on each of the extended arms of the middle housing. (**Supplementary Video 5**)

#### CMOS sub-assembly

The CMOS sub-assembly consists of a custom built fine focus mechanism. The sensor is first attached to a 10mm long cylindrical translational shaft machined out of high-performance plastic (Delrin, Protolabs) via three M1 x 5 mm countersunk cheese headed screws (SFK-M1-5-A2, ACCU) that are inserted through tapped hole provisions provided on the sensor’s PCB board. The opposite face of the shaft holds a thermally inserted M1.2 brass insert (92120A130, McMaster-Carr). A mating hole is machined to create a sliding-fit cylindrical joint with the shaft and is attached with a 1mm thick 304 stainless steel plate using a pair of M1 X 6mm bolts (91800A058, McMaster-Carr) and nuts (HPN-M1-A1, ACCU). The metal plate is pre-fitted with a 10mm long M1.2 bolt (91800A087, McMaster-Carr) and held in place using a nut (90591A312, McMaster-Carr) that is welded to the bolt shaft to form a miniature lead-screw. The shaft is then attached to the miniature leadscrew via the brass insert. Together, the sliding-fit subassembly allows 2mm of bi-directional linear travel of the CMOS sensor within the CMOS enclosure provided on the middle camera housing. Similarly, each of the four CMOS sub-assemblies are attached onto the middle housing using a pair of M1 screws and nuts (ACCU hardware). (**Supplementary Video 5**)

#### CortexCAM final assembly

Once the individual modules are neatly assembled, the top illumination housing is attached to the middle camera housing using four alloy steel socket head screws (93070A278, McMaster-Carr) on the back and front side provisions provided on the scope. Once the assembly process is complete, reusable adhesive putty material is used to cover up the narrow gaps between the LED metal mount and the illumination housing. On the bottom surface, a custom laser cut compressible polyurethane foam piece (86375K141, McMaster-Carr) is attached along the outer edges of the scope aperture to shield the animal’s eyes from light leakage during the experiment. Two square neodymium magnets (B221, K & J Magnetics) are placed into the cavities provided on the implant facing side of the middle housing to attach the Kinematic head-clamp. Two ball point set screws (93339A250, McMaster-Carr) are further inserted into threaded hole provisions provided on the middle housing to fix the position of the kinematic head-clamp relative to the scope. (**Supplementary Video 5**)

#### Electronics and Data Interfacing

We adopted the MiniFAST and miniMscope PCB layouts^18,51^ to fabricate an on-board PCB mounted camera sensor that uses the IMX290LLR, a 1/2.8 optic format back-illuminated CMOS sensor from Sony. The sensor transmits power and data over radio frequency as 8 bit LVDS signals using flexible Coax cable (50 Ohms; CW2040-3650SR, Cooner Wire) that is connected at the sensor end via a custom PCB interface board and at the Data-acquisition side using an SMA coax connector (CONSMA013.062-ND, Digikey) using another interface PCB board. A small amount of UV curing clear resin (Loon UV clear finish, Amazon) is applied to the exposed solder joints to firmly hold the wire connections in place. We use the UCLA Miniscope Data Acquisition (DAQ) for serializing and de-serializing the 8-bit monochromatic pixel data. The firmware and software has been customized to interface with the miniFAST sensor. The DAQ board then sends the data over the USB3.0 interface to the host PC where the camera is controlled using the miniscope camera software version3. This data interfacing framework allows us to use small-format CMOS boards, making it much easier to scale the design to accommodate four imaging modules, each comprising a CMOS sensor and excitation and emission paths, to record neural activity from 4 contiguous fields of view of the brain. A micro-controller (Arduino Uno R3) externally triggers all 4 DAQs to start recording at half the video frame rate (15Hz). The DAQs also registers the system clock time to millisecond precision along with each frame written to memory, allowing us to check for synchronization and calculate frame loss (if any) that might have occurred during synchronized recordings.

The excitation LEDs are connected to the LED driver (LEDD1B, T-Cube LED driver Thorlabs) using 29 gauge multi-stranded wires (9510T213-black, 9510T233-red, McMaster-Carr). The LEDs are operated in the trigger mode to reduce over-exposing the brain to high intensity blue excitation light. A second micro-controller (Arduino Uno 3) is programmed to trigger the LEDs at 15Hz and is synchronized to start with the camera acquisition trigger. Two tactile push buttons are used to initiate and terminate the image acquisition as well as the pulsing of excitation light. (**Supplementary Info. 2a)**

We use a custom commutator (JP41-119-03HW,JINPAT Slip Rings) to interface all four CMOS sensors and eight LED wires to their corresponding DAQs and LED drivers respectively. The Commutator is motorized using Nema stepper motors (NEMA17, Openbuilds Parts Store) and a belt drive system that is tethered to the rotating end^52^. The commutation is controlled wirelessly by a hand-held custom built joystick. An intermediate PCB adaptor, attached with a 14 pin connector head, is used to connect all eight LED wires to the commutator’s rotating side. The coaxial cables are bundled and held together using Kapton tape(KAP22-075, Thorlabs). On the stationery end of the commutator we connect all four coax leads to the DAQ using low loss coaxial connecting cables (RG58, Yotenko). The LED leads are connected to the driver using corresponding LED cables (CAB-LEDD1, Thorlabs). (**Supplementary Info. 2a**)

### Benchtop Optical Characterization

We utilized the 1951 USAF test target (#58-198, Edmund optics) to measure the peak full pitch resolution of a single microscope unit in the CortexCAM assembly. We built a custom 3D printed single camera module to test the resolution. The module consisted of two sub-parts, part one housed the lens and the short pass dichroic filter, and part two housed the CMOS sensor along with the emission filter. The relative positions of all optical components were kept identical to the CortexCAM. The single camera scope was further attached to an xyz translation stage (PT3XYZ stage, Thorlabs) to precisely move the scope in the vertical direction over the test target to bring it to focus. We then calculated the contrast (%) across line pairs from different groups and elements using the below formula:

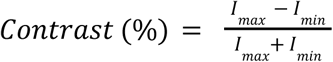

where *I_max_*and *I_min_* represent the maximum and minimum pixel intensity measure across each line pair.

An optical phantom implant prepared by uniformly spreading 10µm green fluorescent microbeads (Lot. 1771416, Elife Technologies) a clear nitrocellulose solution was coated on the underside of the convex polyhedral double glass coverslip implant. The CortexCAM imaged the optical phantom to characterize the resolution and radial distortion in the corners of the implant by measuring the Full Width Half Maximum (FWHM) value for all the micro-beads imaged across a sub-region of the phantom implant window geometry.

In order to test for light uniformity across each individual brain window, we utilized a green fluorescence tape that was cut to fit the shape of each brain window and later attached to the top glass surface. The light intensity variation across each window was measured and quantified using ImageJ and a custom python script.

### Mobile CortexCAM

To operationalize the CortexCAM for neural recordings in freely locomoting mice, we have devised a low-friction sliding and rotating support for the CortexCAM. This additional hardware can be easily built using off-shelf parts and can be adapted to fit a wide variety of behavior arenas. The gantry mechanism was built using four cylindrical guide rails and two cross rails. These rails are mounted onto a custom support frame that is machined from a quarter inch 6061 aluminum plate. The guide rails are placed along the longer edges of the steel frame and placed into custom grooves machined on the support frame and fixed in place using 3D printed end caps. Each of the guide rails are connected to the cross-rails with two low-friction 5 mm linear ball bearings. The two cross rails are mounted onto the guide rails in a 4 x 4 configuration using custom SLA printed mounting brackets (Black resin, Fromlabs) and plastic hardware (92492A703, 93800A300, Mcmaster-Carr). Each of the cross rails is mounted with two miniature 3 mm linear ball bearings. A 3D printed custom carriage that houses an ultra-thin flanged ball bearing (4390N135, McMaster-Carr) is mounted onto the cross rails via the miniature 3 mm linear ball bearings using removable 3D printed mounting caps and light weight plastic fasters. This design prevents binding of the rails across the entire workspace (800mm X 500mm) because of the even distribution of loads on each linear ball bearing preventing overload and the compliance in using end-mounted cylindrical rails. The CortexCAM can be attached to the custom carriage by press-fitting the flange connector of the scope into the inner-ring of the flanged ball bearing housed within the carriage. For additional fixation a circular locking collar (CNC machined from Delrin, Protolabs) is attached to the outer ring of the flanged bearing via three brass screws (96741A120, McMaster-Carr) that can be driven into tapped hole provisions provided on the flange connector of the CortexCAM. Once assembled, the platform allows the end-effector, which in our case is the CortexCAM, to translate freely in the x-y plane and rotate about the yaw axis. The final assembly is attached onto an elevated stage built by vertically stacking three T slotted framing rails (47065T101, McMaster-Carr), aligned lengthwise along the two longer edges of the support frame, using compatible hardware and fixing brackets.

### Surgery

All animal experiments were approved (protocol no. 2202-39800A) by the University of Minnesota Institutional Animal Care and Use Committee. Wildtype C57/B6 mice (n = 3) were used for initial testing of locomotion and clamping mechanisms. Double transgenic mice (n = 3 mice) derived from cross-breeding Cux2-Cre-ERT2 (RRID:MMRRC-032779-MU, MMRRC) and Ai162-GCaMP6s^53^ were utilized for the Ca2+ imaging experiments. Mice were housed in a 14-hour light/10-hour dark cycle in rooms maintained at 20 - 23 °C and 30 - 70 % relative humidity. Double transgenic mice were injected with tamoxifen (75 mg/kg) for 5 days before surgery to induce sparse expression of GCaMP in cortex layers 2 and 3 pyramidal neurons^3^. Surgical methodologies from our previous studies^13,54,3,18^ were adapted for implanting the convex-polyhedral coverslip implants. Briefly, ∼60 minutes before the surgery, mice were administered sustained-release buprenorphine (1 mg/kg; Buprenorphine SR-LAB, Wedgewood Connect) and dexamethasone (2mg/kg) for analgesic and preventing neuroinflammatory response respectively

Mice were first anesthetized in an induction chamber with 1-5% isoflurane in oxygen. An eye ointment (Puralube, Dechra Veterinary Products) was applied to the eyes. The scalp was shaved and the mice were transferred to a stereotaxic (900LS, Kopf). After fixation to the stereotax, the scalp was sterilized by applying Betadine and 70% ethanol solution using a cotton tipped applicator. Following this the scalp was removed using surgical scissors. The 3D frame of the convex polyhedral cranial window implant was placed on the skull. The cranial window was positioned symmetrically about the medial longitudinal fissure and the centroid of the implant was positioned approximately 4mm anterior to the Bregma. A curved dental probe was used to score the outer edges. A dental drill was then used to remove the bone along the score perimeter to open the craniotomy. Once the bone island was fully detached along the edges, we carefully removed it using surgical tweezers and cotton tipped applicators. The exposed brain surface was flushed with sterile saline and immediately covered using a gel foam soaked in sterile saline until the bleeding from the excised bone stopped. The gel foam was removed and the cranial window was gently lowered onto the surface of the brain and a slight amount of pressure was applied using an applicator over the implant. A surgical adhesive (Vetbond, 3M) was applied along the outer 3D printed bezel of the implant to fix it onto the skull while ensuring no gaps between the implant surface and the skull were present. An additional layer of dental cement (Metabond, Parkell Inc.) was applied along the edges for further adhesion. After the dental cement cured, a titanium headplate was attached to the cranial implant using 0-80 screws. Post surgery, the mice were moved onto a heating pad to recover and later transferred into a clean cage until they were ambulatory. All animals were allowed to recover for 6-7 days before proceeding with the imaging or behavior experiments.

### Head-fixation and habituation

A custom head-fixation stage was built using half inch optical head posts (TR4, TR6, Thorlabs), post holders (PH4, Thorlabs), and compatible construction accessories (RA90, SWC, Thorlabs). The CortexCAM was attached to the head-fixation station using a custom designed stainless-steel adaptor that was mounted onto a 3D printed cross-beam. Coarse adjustment of the height of the head-dock was performed by loosening the fixing bolts provided on either side of the cross-beam. A 3D printed treadmill with an integrated quadrature optical encoder (B07MX1SYXB, Amazon) was incorporated underneath the CortexCAM. Two behavior cameras (IMX273 U3-162SM-CS, Blackfly USB 3.1 FLIR) were set up to capture the front and side profile of the mouse respectively. An external trigger was provided by a micro-controller (Arduino Uno3) to initialize frame acquisition for the CortexCAM and the behavior cameras. The mice were first handled for 3 days followed by 3 days of head-fixation to acclimatize to the treadmill and the head-docking mechanism (**Supplementary Video 2**).

### Freely locomoting mice behavior testing on the figure 8 maze

We built a custom automated figure 8 maze arena measuring 80 x 50 cm under the mobile CortexCAM setup. The arena consisted of 3D printed walls with a 15 mm recessed bay feature that would allow the mice to fully explore the arena without any hindrance while tethered to the mobile CortexCAM. The floor was made from a custom cut ¼ inch acrylic sheet overlaid with 6 mm thick rubber foam sheet (EVA foam, Amazon). Three IR break-beam sensors(1528-2526-ND, DigiKey) were incorporated in each of the arms of the 8 maze arena to detect animal motion. Servo actuated swing doors (SG90, 9g Mirco-servos, Amazon) were placed at the entrance at exit of the central decision arm of the maze. The arena was supported by half inch custom machined cylindrical metal posts with swivel levelling mounts (611K46, McMaster) to accommodate any z height adjustments required while tethering animals to the mobile CortexCAM system. An enclosure was built to house the figure 8 maze arena and the mobile CortexCAM platform using 80/20 T slotted aluminium rails and ¼ inch medium density fiber board sheets (McmasterCarr). Additionally, the inside walls of the enclosure were sound-proofed using acoustic foam panels (B08QN6B2FD, Amazon). The arena was dimly lit using incandescent string lights (B07TWV1WGJ, Amazon), allowing the animal to navigate the space. We also had provisions to increase illumination within the enclosure using red led strip lights and IR illumination light sources (B07MX63RX3, Amazon). The commutator was mounted using custom 3D printed brackets and a t-slotted cross-rail attached to the enclosure roof rails. A top mounted behavior camera (IMX273 U3-162SM-CS, Blackfly USB3.1 FLIR) (**Supplementary Info. 2b)** was used to capture the animal’s movement. A second camera was used to capture the side perspective view of the animal moving in the arena. The mice were head-fixed onto the mobile CortexCAM by fixing all three degrees of freedom. We used rubber tipped paper clips to arrest the x or y translation of the carriage along the cylindrical rails and a custom 3D printed key that locked the rotation of the CortexCAM about the yaw axis. The animal was allowed to move on a horizontal mouse wheel that was integrated with the figure eight maze arena while the experimenter docked the animal to the mobile CortexCAM station. The experimenter then tightened the set screws to complete the tethering procedure. Additionally, the experimenter also made fine focus adjustments to the scope for imaging before each recording session. Once the animal was docked onto the platform, the paper clips on the guide rails were removed and the wheel motion was arrested, allowing the animal to self-initiate movement into the figure eight maze arena. Once the animal entered the arena, the remaining two clips on the cross rails and the 3D printed locking key were removed sequentially, allowing the animal unrestrained movement in the arena along the x, y and yaw directions while being tethered to the CortexCAM platform (**Supplementary Video 3)**.

### Sociability test

A three chamber sociability arena^55^ was adapted with shallow recessed walls to fit under the mobile CortexCAM. The animals were first habituated to the arena by placing them in the central chamber with empty side chambers for 10 minutes. During the actual experiment trial the animals were tethered to the mobile CortexCAM platform and allowed to self initiate movement into the 3 chambered arena using similar protocol as followed for the figure 8 maze experiment. Mice were first restricted to exploring the central chamber of the behavior arena for the first 10 minutes. This was followed by allowing the animal to explore all three chambers for another 10 minutes by removing the barricades that were placed at the entrances of the two peripheral chambers. Next, a juvenile target mouse was placed inside a cylindrical wired mouse cage in one of the side chambers. The wired cages were modified to have a dome shaped ceiling to allow the test mouse, tethered to CortexCAM, to approach and interact with the conspecific without any hindrances. A novel inanimate object was placed inside an identical cage on the opposite chamber. The test lasted 20 minutes where we acquired neural recordings from the test mouse using the CortexCAM and simultaneously tracked the social interactions between the test and target mice using two behavior cameras (**Supplementary Video 4)**.

### Behavior video acquisition and analyses

Each frame captured in the behavioral imaging cameras was triggered using a TTL pulse signal generated by the neuroimaging data acquisition system (Miniscope DAQ V 3.2, LABmaker) at the onset of a frame recorded on the CortexCAM. This allowed us to synchronize both the behavior and the multiple CMOS sensors in the CortexCAM. Data from the behavior cameras were acquired, on a separate portable PC (Mini S13 Pro, Beelink), using their native camera software (Spinnaker SDK, Teledyne) to offload memory utilization on the parent computer that is used for recording the neural data.

### Pose estimation and head-direction measurements

Markerless pose estimation (Deeplabcut, V 3.0.0)^40^ was used to detect up to four key-points to estimate the position, speed and the heading direction of the mouse tethered to the mobile CortexCAM. The rigid fixation of the mouse implant relative to the CortexCAM ensured that we consistently tracked the head and neck position of the mouse without any perspective and depth ambiguities in the pose estimation. The head direction was estimated by drawing a vector between the two key-points used for identifying the head and neck positions of the animal. The raw position data obtained from markerless tracking was further filtered using a custom python code to remove outliers and replace predicted key-points with low confidence levels (likelihood <0.3) with the mean of the position data from the neighboring frames.

### Ca^2+^ imaging data analysis

Motion correction, cell registration and cell trace extraction from our calcium video datasets was performed using Suite2p^36^. We optimized the default 1P imaging settings to best fit our dataset. The decay time (*Tau*) of 1.5 s and the frame rate of 15Hz were updated in the main settings to match our experimental parameters. We also set the threshold scaling factor to 0.25 to improve cell detection probability^36^. We enforced the spatial scale of ROI to be 6 pixels as identified in our dataset. The ROI detection algorithm was first executed in an iterative loop with maximum iteration set at 20. We then manually curated the dataset to classify ROIs as cells and the rest as not cells by first checking if the ROI was spatially distinct from their neighboring cells and was well separated from blood vessels. Then we checked and verified if the corresponding trace activity had a typical Calcium signature with instant peaks and gradual decay above the neuropil activity levels. We also used the aspect ratio and cell class classification tools provided in Suite2p GUI to estimate the quality of the cell’s footprint. After manual curation, we performed neuropil subtraction on the raw traces as done in previous studies^56^. The DFF was calculated as the ratio of fluorescence increase over baseline fluorescence (F0), where F0 was approximated to be the 8^th^ percentile of the cell’s neuropil corrected temporal trace data^57^. The deconvolved traces were extracted from the DFF trace data by specifying a constant decay timescale of 1.5s for GCamP6s^58^.

### Composite brain map and alignment to the Allen Brain Atlas

For each FOV of the CortexCAM, an affine transformation was estimated by registering a reference frame to a fluorescence widefield image of the whole cranial window implant obtained using a benchtop microscope (Leica MZ 10F). We then applied rigid transformation on the widefield image to align it with a 2D projection of the Allen Common Coordinate Framework v3(CCF)^59,60^ as done in past work on mesoscopic calcium imaging studies^31,61^. The transformations were applied to map the pixel coordinates of each neuron centroid, obtained after cell extraction using Suite2p, to the global frame of reference on the widefield image that is in turn aligned with the Allen CCF. Each ROI location was masked using the anatomical regions defined by the Allen brain atlas.

### Heading direction encoding analysis

The heading direction analysis was performed based on the methods outlined in recent heading direction studies conducted in freely behaving mice experiments^62^. The cellular calcium traces, their corresponding anatomical mappings on the Allen brain atlas, centroid location of each ROIs in the global frame of reference, the instantaneous mouse position, time stamps of frame capture, and heading angle of the mouse were combined to form a unified pandas dataframe using a custom python script. The deconvolved spike train data was further smoothed using a moving average filter of width 5 frames (∼330 ms) to give us the firing activity of each neuron. We further partitioned the azimuth plane to 10 degree angle bins and assigned the instantaneous heading angle of the mouse to their corresponding angle bin. The tuning curve was calculated as the occupancy normalized firing activity of the cell within each angle bin. The tuning curves are then circularly smoothed using a moving average filter of width 50°. The preferred firing direction (PFD) of the cell is defined as the angle bin corresponding to peak tuning strength value. A stimulus signal is then derived for each angle bin using the following equation:

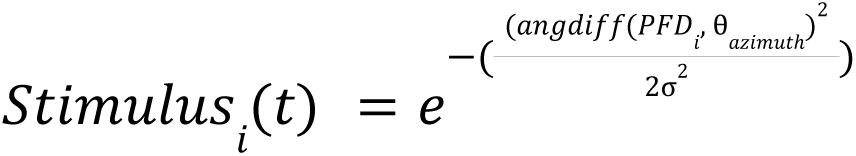

Where θ_𝑎𝑧𝑖𝑚𝑢𝑡ℎ_ is the estimated heading angle bin and σ is the standard deviation for the gaussian kernel (σ = 2). The subscript 𝑖 ranges from 1 to 36, representing each angle bin. The angular difference function, 𝑎𝑛𝑔𝑑𝑖𝑓𝑓(𝑎, 𝑏) estimates the circular difference between 𝑎 and 𝑏 wrapped between [− π, π].

We then calculated Pearson’s correlation of the stimulus signal and the normalized spiking activity for each cell. A null distribution of the deconvolved DFF traces were synthesized by applying 1000 random shuffles of the spiking activity for each cell. This was utilized to determine the threshold for classifying cells as highly tuned to heading direction vs less tuned to heading direction through an iterative process. This step is only performed once initially to determine the threshold values for a particular session. The cells that are classified as highly tuned to heading direction are further filtered out from the parent dataframe and sorted based on heading direction. The candidate cells for each angle bin are then selected by estimating the least circular variance of their corresponding tuning strength curves.

### Statistical testing

A Friedman test with a Tukey post hoc test was performed to examine differences in the distance moved, speed metrics, number of trails and the percentage of correct choices across five experimental conditions, (7 g dummy scope, 28 g dummy scope, 52 g dummy scope, tethered to mobile CortexCAM, untethered and freely behaving), with three subjects as shown in **Supplementary Info. 3 a-d**. Friedman’s test was chosen due to the non-parametric nature of the given sample size of three subjects. The test has been used previously in similar experiments with smaller sample sizes^25^. We randomized the order of behavior trials while testing the influence of different payloads on the mouse tethered to CortexCAM.

## SUPPLEMENTARY VIDEOS

**Supplementary Video 1:** Comparison of FOV of CortexCAM with other setups

**Supplementary Video 2:** Headfixed imaging using the CortexCAM with multiperspective behavior imaging

**Supplementary Video 3:** Behavior in mobile CortexCAM with imaging

**Supplementary video 4:** Sociability test behavior with imaging

**Supplementary Video 5:** Assembly of the CortexCAM

## DATA AVAILABILITY STATEMENT

All data needed to evaluate the conclusions in the paper are present in the paper and/or the Supplementary Materials. Additional datasets required for generating the figures are accessible through Zenodo: https://doi.org/10.5281/zenodo.18496840

## CODE AVAILABILITY STATEMENT

All codesets used for analysis of the neurobehavioral data are made available through the github repository: https://github.com/bsbrl/CortexCAM

## ACKNOWLEDGEMENTS

We acknowledge the staff at Research Animal Resources, University of Minnesota, for animal care and housing. The mouse strain used for this research project was obtained from the Mutant Mouse Resource and Research Center (MMRRC) at University of Missouri, an NIH-funded strain repository, and was donated to the MMRRC by Ulrich Mueller, Ph.D., The Scripps Research Institute. We thank Peter Ness for custom machining support. We thank James Hope for the fruitful discussions while developing the docking mechanism. We thank Hamza Beder for consulting on custom PCB design. We thank Farnoosh Dadashi for assistance with animal injections. We also thank Janaki Riji Nair for helping with the statistical analysis for this work.

## AUTHOR CONTRIBUTIONS

A.C., Z. V. and S.K. conceptualized the CortexCAM. A.C and Z.V. designed the optical architecture. A.C performed the bench top characterization for the CortexCAM. A.C. and Z.V. designed and built the mobile CortexCAM platform. A.C developed the CortexCAM imaging hardware and software. A.C., J.H., S.F., E.K. and K.S. performed animal surgery procedures and injections. A.C., S.F., Z.V., and J.H. worked on implant development. A.C., J.J. and I.O. developed the PCB design for imaging sensors. A.C., J.J. and D.S. developed the multi camera imaging firmware. A.C. and I.O. developed the motorized commutator. A.C., V.K. and K.S. conducted animal behavior training. A.C, V.K. and K.S. performed the imaging experiments. A.C. processed the raw experimental data. A.C., D.S. and S.K. performed the data analysis. A.C., Z.V. and D.S. developed the alternating choice behavior assay. A.C., S.P., S.S., and M.L. developed the sociability task behavior assay. A.C. and S.K. visualized data. A.C and S.K. wrote the original manuscript. Every author reviewed and editing the manuscript. A.C. and S.K. supervised all aspects of the work. R.H. and S.K. acquired funding.

## Supplementary Information

**Supplementary Information 1:**
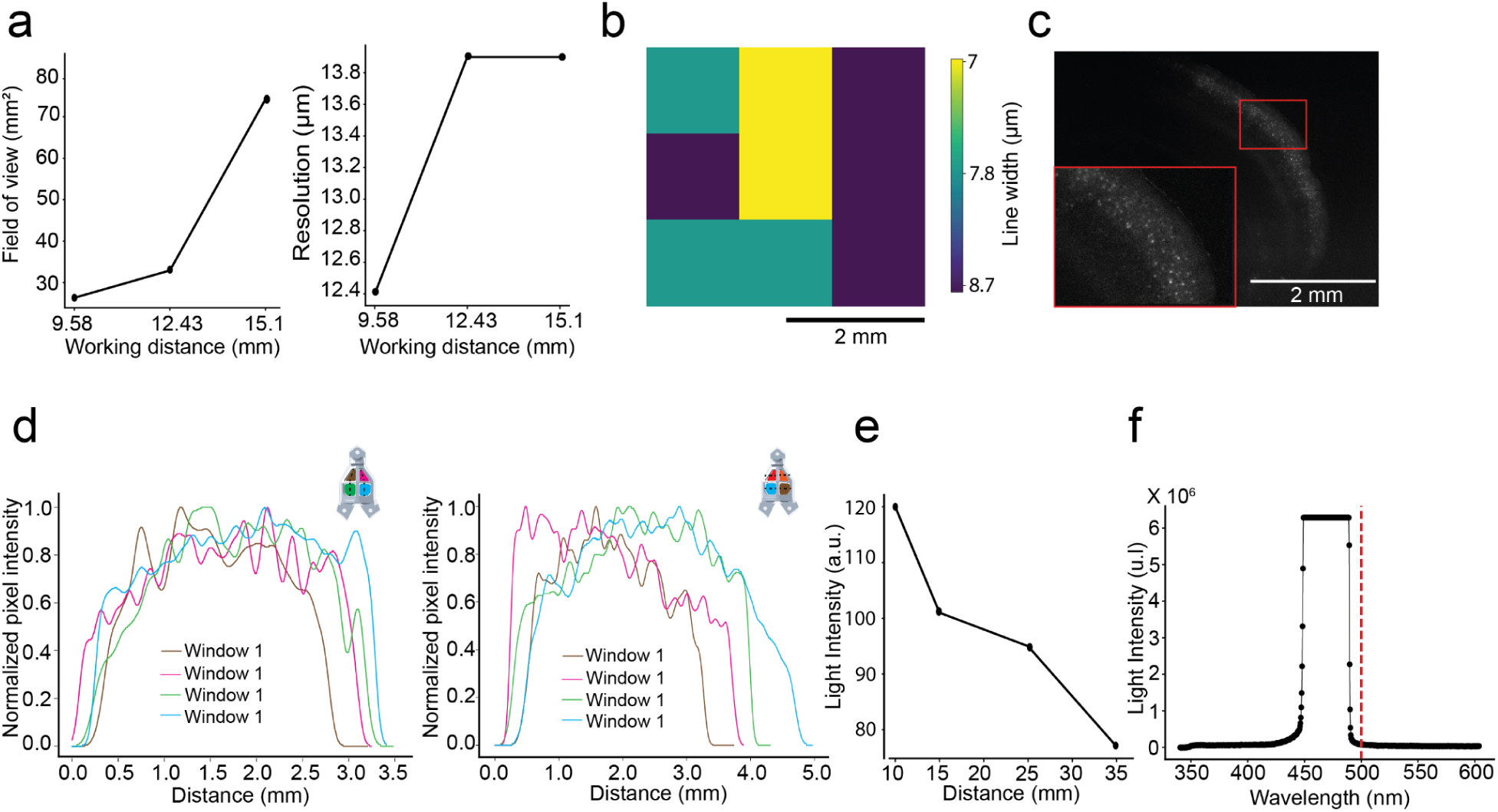
Additional benchtop optical characterization results for CortexCAM. a) Changes in FOV and resolution for the CortexCAM as a function of varying working distances obtained using a single camera module replica of CortexCAM b) A Pseudo color heat map representing the variation in resolution across the field of view imaged using a single camera module replica of CortexCAM. c) A 400μm coronal brain slice from a Cux2-CRE-ERT2 x Ai162 (TIT2L-GC6s-ICL-tTA2) imaged using a single camera module replica of the CortexCAM. The inset shows a zoomed in view of a sub-region that shows individual cell bodies present in layers 2 and 3 of the brain slice. d) Light uniformity measurements made using a phantom implant, attached with a fluorescent tape on the bottom surface of the implant, across all four FOVs along the vertical and horizontal axes as shown in the respective inset figures. e) Change in light intensity of the collimated light source module as a function of distance from the output aperture. f) The measured light power spectrum for the collimated excitation light source with the red dashed line showing a cut off below 500nm.

**Supplementary Information 2:**
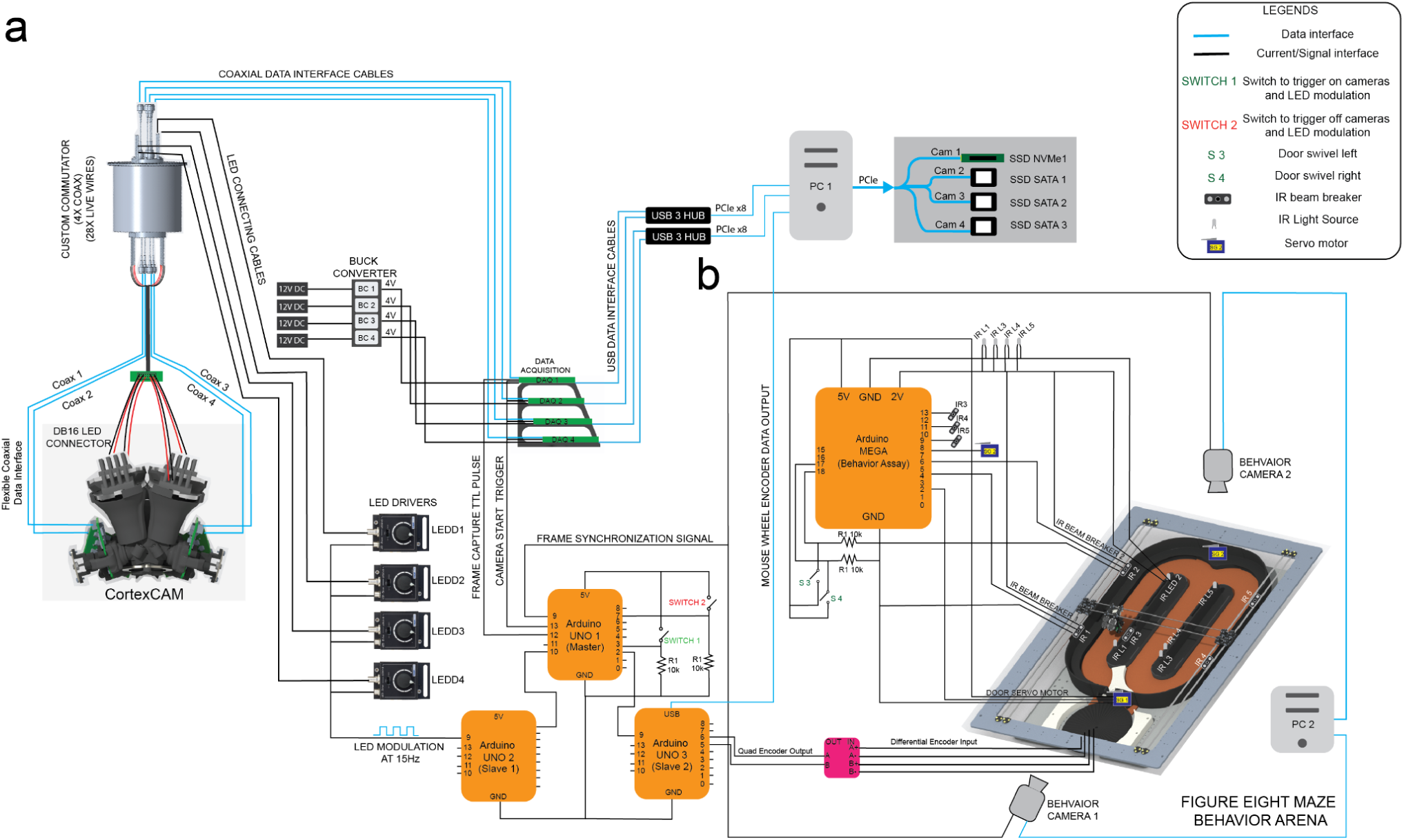
Mobile CortexCAM electrical wiring diagram. a) The wiring schematic for the CortexCAM device with blue connecting lines showing the data streaming, acquisition and storage connections and the black lines indicating the hardware automation, camera synchronization, LED modulation connections. b) The wiring schematic for the automated figure eight maze behavior assay.

**Supplementary Information 3:**
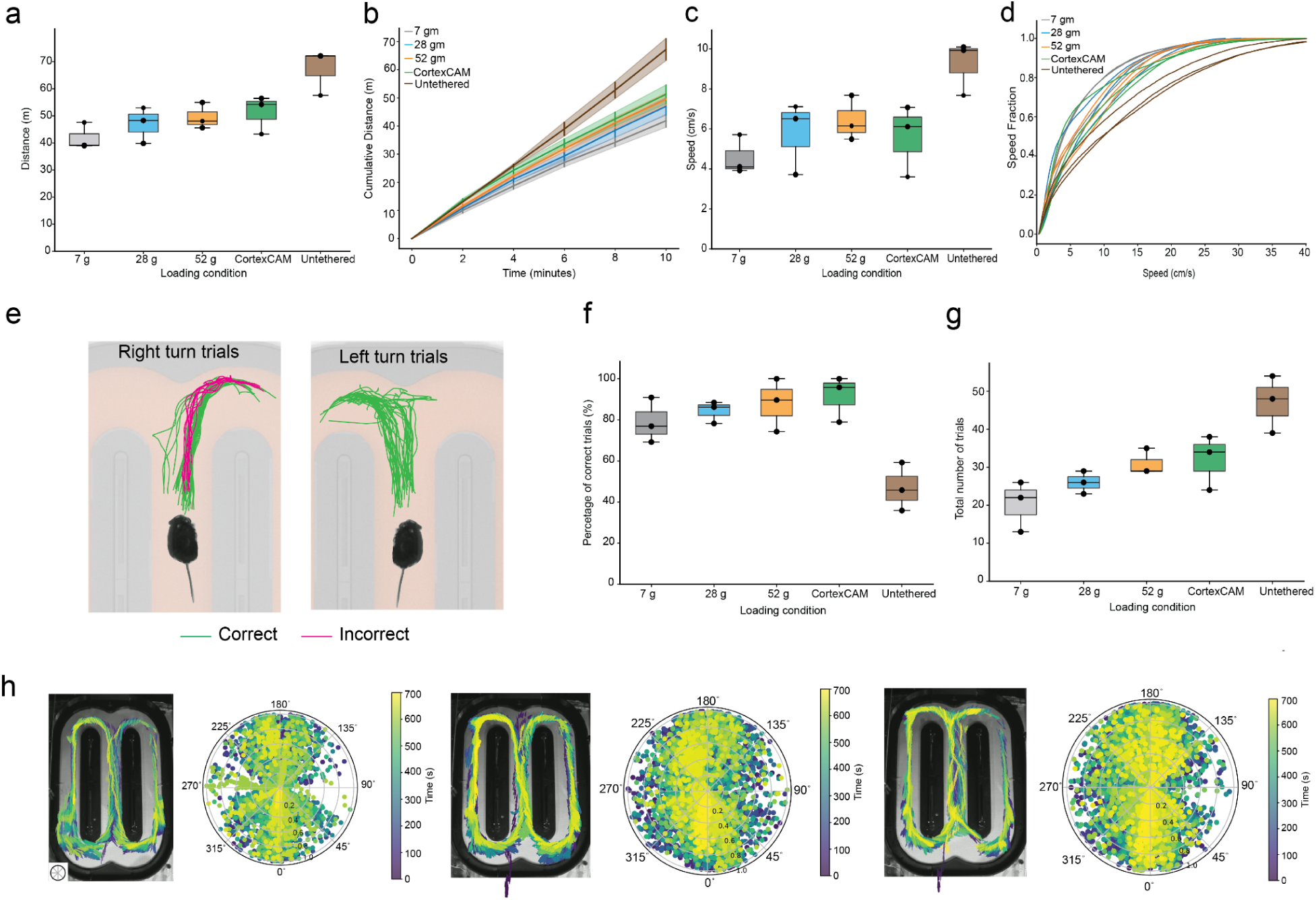
Behavior data analysis of mice performing figure eight maze task. a) Distance moved by mice (n=3) across different loading conditions (Friedman test indicated that there is no statistically significant difference in distance moved by the subject between the conditions, χ^2^ (4, *N* = 3) = 8.27, *p* = 0.08). b) Cumulative distance moved by the mice (n=3) across 10 minute trial durations during different loading conditions. c) Median speed distribution of mice (n=3) while performing a figure eight maze task during different loading conditions. (Friedman test showed no significant difference between conditions for speed of the subject, χ^2^ (4, *N* = 3) = 7.73, *p* = 0.10). d) The cumulative speed distribution of the mouse across different loading conditions while performing the figure eight maze task. e) Trajectory of mice across all trials at the decision zone during the figure eight maze task. The correct trials are marked in green and the incorrect trials are naked as red. f) Percentage of correct alternations by the mice (n=3) across different loading conditions. (The Friedman test showed statistical differences between conditions for the percentage of correct choices, χ^2^ (4, *N* = 3) = 10.37, *p* = 0.03). g) Total number of trials completed by mice (n=3) across different loading conditions.(Friedman test for the total number of trials performed during a 10 minute session showed no statistically significant differences between the conditions, χ^2^ (4, *N* = 3) = 8.61, *p* = 0.07). h) The trajectory and heading angle distribution of all three mice during a single trial performing the figure eight maze alternating choice task while tethered to the fully assembled CortexCAM.

